# An attempt to replicate a dissociation between syntax and semantics during sentence comprehension reported by Dapretto & Bookheimer (1999, *Neuron*)

**DOI:** 10.1101/110791

**Authors:** Matthew Siegelman, Zachary Mineroff, Idan Blank, Evelina Fedorenko

## Abstract

Does processing the meanings of individual words *vs*. assembling words into phrases and sentences rely on distinct pools of cognitive and neural resources? Many have argued for such a dissociation, although the field is lacking a consensus on which brain region(s) support lexico-semantic *vs*. syntactic processing. Although some have also argued against such a dissociation, the dominant view in the field remains that distinct brain regions support these two fundamental components of language. One of the earlier and most cited pieces of evidence in favor of this dissociation comes from a paper by Dapretto & Bookheimer (1999, Neuron; DB). Using a sentence meaning comparison task, DB observed two distinct peaks within the left inferior frontal gyrus (LIFG): one more active when comparisons relied on lexico-semantic cues, and another – when they instead relied on syntactic cues. Although the paper has been highly cited over the years, no attempt has been made, to our knowledge, to replicate the original finding. We here report an fMRI study that attempts to do so. Using a combination of three approaches – whole-brain, group-level ROIs, and individual functional ROIs – we fail to replicate the originally reported dissociation. In particular, parts of the LIFG respond reliably more strongly during lexico-semantic than syntactic processing, but no part of the LIFG (including in the region defined around the peak reported by DB) shows the opposite response pattern. We hypothesize that the original result was a false positive, possibly driven by one participant or item given the use of a fixed-effects analysis and a small number of items (8 per condition) and participants (*n*=8).

## 1. Introduction

Sentence comprehension requires us to understand both the meanings of individual words and the dependency structure between them (conveyed via word order and/or functional morphology), which determines the propositional content of the sentence (i.e., who is doing what to whom). Whether these two components of sentence comprehension rely on distinct pools of cognitive and neural resources has been long debated.

A number of manipulations have been used in an attempt to selectively target lexico-semantic *vs.* syntactic/structural processing, including (i) linguistically degraded materials like lists of unconnected words that require lexical-level understanding but not putting words together into more complex representations, *vs.* “Jabberwocky” sentences that contain a coarse-level representation of the dependency structure but not lexical meanings (e.g., Friederici et al., 2000; Humphries et al., 2006; Fedorenko et al., 2010); (ii) violations of lexico-semantic *vs.* syntactic expectations (e.g., Embick et al., 2000; Cooke et al., 2006; Friederici et al., 2010; Herrmann et al., 2012), an approach that proved fruitful in ERP investigations of language (e.g., Kutas & Hillyard, 1980; Kutas & Federmeier, 2011; Osterhout & Holcomb, 1992; Hagoort et al., 1993); and (iii) adaptation to lexico-semantic content *vs.* syntactic structure (e.g., Noppeney & Price, 2004; Santi & Grodzinsky, 2010; Menenti et al., 2012; Segaert et al., 2012). These numerous studies have produced a complicated empirical picture with some contradictory results. Nevertheless, the dominant view in the field remains that lexico-semantic and syntactic processing rely on distinct brain regions, and, by extension, are cognitively separable (cf. Bates & Goodman, 1997; Fedorenko et al., 2012a; Blank et al., 2016; Wilson & Bautista, 2016).

One of the most cited studies that argues for a dissociation between semantic and syntactic processing was conducted by Dapretto and Bookheimer (henceforth: DB) and published in Neuron in 1999. The study used an original manipulation where participants compared meanings across pairs of sentences, with the relevant information for deciding whether the two sentences had the same or different meanings being either lexico-semantic or syntactic. In the former condition, one sentence differed from another in a single word, replaced by either a synonym (same) or a non-synonym (different); in the latter, the two sentences differed in structure, as in the Active/Passive alternation in which the thematic roles were either kept constant (same) or were switched (different) (see sample items in section 2.2.1). The key result was a dissociation between the Semantic and Syntactic conditions observed in the left inferior frontal gyrus (LIFG), with two nearby but distinct peaks revealed by the *Semantic>Syntactic* contrast and *Syntactic>Semantic* contrast, respectively. This result, the authors argued, provided “unequivocal evidence that these functions [semantic and syntactic processing; *SMBF*] are […] subserved by distinct cortical areas”.

DB's study has been cited 632 times (as of February 21, 2017; Google Scholar), and the pattern of citations over the years (Figure 1) suggests that it is still being used by researchers as evidence for distinct brain regions supporting semantic *vs*. syntactic processing (citations per year, from 2000 to 2016: *M*=36.6, *SD*=21; *Median*=34, *Interquartile range*=15.2).

**Figure 1.**
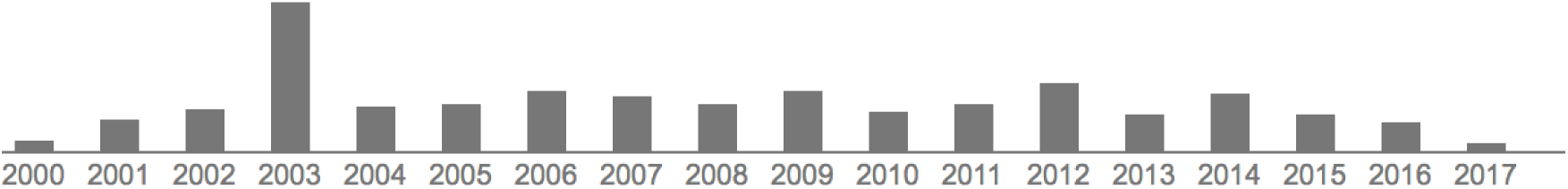
(from Google Scholar) The citations for DB (1999) distributed over the years (total *n* = 632 as of February 21, 2017).

And yet, it appears that this study has never been replicated either by one of the original author’s labs, or by any other research group. Given (i) the study’s impact on the field, combined with (ii) recent studies that have argued for overlap between lexico-semantic and syntactic processing across the fronto-temporal language network (e.g., Blank et al., 2016; Wilson & Bautista, 2016; Fedorenko et al., 2017, in prep.), and (iii) current emphasis on reproducibility in the fields of psychology (e.g., Ioannidis, 2005; Simmons et al., 2011; Button et al., 2013; Ioannidis et al., 2014) and cognitive neuroscience (e.g.,Poldrack et al., 2017), we here attempted to replicate DB's results.

## 2 Methods

### 2.1. Participants

Fifteen individuals between the ages 20-30 (*M*=25.3, *SD*=4.1; 5 females), all native speakers of English, participated for payment. Fourteen of the fifteen participants were right-handed (as determined by the Edinburgh handedness inventory; Oldfield, 1971), but all fifteen showed typical, left-lateralized, language activations (as assessed with an independent language “localizer” task conducted in the same session; Fedorenko et al., 2010). All participants had normal hearing and vision, and no history of neurological illness or language impairment. Participants gave informed consent in accordance with the requirements of MIT’s Committee on the Use of Humans as Experimental Subjects (COUHES).

### 2.2. Design, materials, and procedure

Each participant completed the critical task, as well as one or more additional tasks for unrelated studies. The entire scanning session lasted approximately 2 hours.

#### 2.2.1. Design and materials

The basic design was the same as in the DB study: participants were presented with pairs of sentences and asked to decide whether they mean roughly the same thing. The critical manipulation was whether the sentences in the pair differed in one of the words (the Semantic condition) or in the structure / word order (the Syntactic condition). In particular, in the Semantic condition, one sentence differed from the other in a single word, replaced by either a synonym (same meaning) or a non-synonym (different meaning), as in (1a) below. In the Syntactic condition, the two sentences were either syntactic alternations differing in both structure and word order (same meaning), or in only structure / only word order (different meaning), as in (1b):

(1a) Semantic condition
  Same: *Anna invited the composer. / Anna invited the songwriter.*
  Different: *Anna invited the composer. / Anna invited the translator.*
(1b) Syntactic condition
  Same: *Anna invited the composer. / The composer was invited by Anna.*
  Different: *Anna invited the composer. / The composer invited Anna.*

The materials consisted of 80 items (sentence pairs). Forty items used the Active / Passive constructions (as in DB), and forty – the Double Object (DO) / Prepositional Phrase Object (PP) constructions. Each item had four versions, as in (1a-b), for a total of 320 trials. The full set of materials is available at https://evlab.mit.edu/papers/Siegelman_bioarxiv.

The 320 trials were divided into four experimental lists (80 trials each, 40 trials per condition) following a Latin Square Design so that each list contained only one version of any given item. Any given participant saw the materials from just one experimental list, and each list was seen by 3-4 participants.

A number of features varied and were balanced across the materials. First, the construction was always the *same* across the two sentences in the Semantic condition (balanced between active and passive for the Active/Passive trials, and between DO and PP for the DO/PP trials). However, in the Syntactic condition, the construction was always *different* in the Same-meaning trials because this is how the propositional meaning was preserved (again, balanced between active and passive for the Active/Passive trials, and between DO and PP for the DO/PP trials). For the Different-meaning trials, the construction could either be the same or different, as follows:

(2a) Syntactic condition – Different meaning, Active/Passive:

Same construction: *Anna invited the composer. / The composer invited Anna.*
Different constructions: *Elizabeth disliked the proprietor. / Elizabeth was disliked by the proprietor.*
(2b) Syntactic condition – Different meaning, DO/PP:

Same construction: *Amanda lent the cook some money. / The cook lent Amanda some money.*
Different constructions: *Brenda read the expert a passage. / The expert read a passage to Brenda.*

For trials where the constructions differed between the two sentences in a pair, we balanced whether the first sentence was active *vs.* passive (for the Active/Passive trials), or whether it was DO *vs.* PP (for the DO/PP trials).

The second feature that varied across the materials was the whether the first-mentioned noun was a name or an occupation noun. All sentences (in both Active/Passive and DO/PP constructions) contained one instance of each, with order of presentation balanced across items.

And third, for the Semantic condition, we varied how exactly the words in the second sentence in the pair differed from the words in the first. (This does not apply to the Syntactic condition trials because the content words are identical across the two sentences within each pair.) In particular, for the Active/Passive trials, either the occupation noun or the verb could be replaced (by a synonym or a word with a different meaning); and for the DO/PP trials, either the occupation noun or the direct object (inanimate) noun could be replaced.

#### 2.2.2. Procedure

An event-related design was used. Each event (trial) consisted of an initial 300ms fixation, 2,000ms presentation of the first sentence (presented all at once), 200ms inter-sentence interval, 2,000ms presentation of the second sentence, and a 1,500ms window for participants to respond (by pressing one of two buttons on a button box), for a total of 6s. The 80 trials in a list were divided into two runs, with each run consisting of 40 trials and an additional 120s of inter-trial fixation, for a total run duration of 360s (6min). Each participant performed two runs. The optseq2 algorithm (Dale, 1999) was used to create condition orderings and to distribute fixation among the trials so as to optimize our ability to de-convolve neural responses to individual trials. Eight orders were created, and order varied across runs and participants.

#### 2.2.3. A summary of the key differences between our study and DB's study

Table 1 provides a summary of the key differences between the studies. The key improvements in the design of the current study concern statistical power, both with respect to the number of participants tested (almost twice as many participants), and the experimental materials (five times as many trials per condition). Note that even though the original study used blocked design, which is generally more powerful than event-related designs, it consisted only of one block per condition; thus, the current study is likely to be vastly more powerful in spite of the use of an event-related design (with 40 events per condition). We also deviated in one of the constructions used. Although we adopted the Active/Passive alternation, we replaced the Locative Prepositional Phrase alternation (e.g., *The pool is behind the gate. / Behind the gate is the pool.*) with a more commonly used Double Object / Prepositional Phrase Object (DO/PP) alternation (e.g., Allen et al., 2012; Gibson et al., 2013). The reason we chose not to use the Locative alternation from the original study is that fronted locative prepositional phrases (locative inversion) are rare in natural language (e.g., Gibson et al., 2013). Finally, we opted for the use of the visual presentation (cf. auditory presentation used by DB). Abundant prior evidence suggests that high-level language processing brain regions, including those in the frontal lobe, are robust to presentation modality manipulations (e.g., Fedorenko et al., 2011; Braze et al., 2011; Vagharchakian et al., 2012; Scott et al., 2016). Thus, the use of a different modality is not expected to matter.

**Table 1:** A summary of the key differences between the two studies.

|  | **Current study** | **DB (1999)** |
| --- | --- | --- |
| Participants | *n*=15 | *n*=8 |
| fMRI design | Event-related | Blocked |
| Materials | 40 pairs of sentences per condition, 40 events per condition | 8 pairs of sentences per condition, 1 block per condition |
| Constructions used | Active/Passive alternation, Double Object/ Prepositional Phrase Object alternation | Active/Passive alternation, Locative Prepositional Phrase alternation |
| Presentation | Visual | Auditory |

### 2.3. fMRI data acquisition and preprocessing

Structural and functional data were collected on the whole-body 3-Tesla Siemens Trio scanner with a 32-channel head coil at the Athinoula A. Martinos Imaging Center at the McGovern Institute for Brain Research at MIT. T1-weighted structural images were collected in 128 axial slices with 1mm isotropic voxels (TR = 2530ms, TE = 3.48ms). Functional, blood oxygenation level dependent (BOLD) data were acquired using an EPI sequence (with a 90° flip angle and using GRAPPA with an acceleration factor of 2), with the following acquisition parameters: thirty-one 4mm thick near-axial slices, acquired in an interleaved order with a 10% distance factor; 2.1mm×2.1mm in-plane resolution; field of view of 200mm in the phase encoding anterior to posterior (A>P) direction; matrix size of 96mm×96mm; TR of 2000ms; and TE of 30ms. Prospective acquisition correction (Thesen et al., 2000) was used to adjust the positions of the gradients based on the participant's motion one TR back. The first 10s of each run were excluded to allow for steady-state magnetization.

MRI data were analyzed using SPM5 and custom MATLAB scripts. Each participant’s data were motion corrected and then normalized into a common brain space (the Montreal Neurological Institute Brain Template; MNI) and resampled into 2mm isotropic voxels. The data were then smoothed with a 4mm FWHM Gaussian filter and high-pass filtered at 200s. The effects of experimental conditions were estimated using a General Linear Model (GLM), with each experimental condition modeled by a boxcar function (corresponding to an event) convolved with the canonical hemodynamic response function (HRF).

### 2.4. Data analysis

We performed three analyses to assess whether the dissociation between syntactic and semantic processing, as reported by DB, holds in the current dataset.

#### 2.4.1. Traditional, group-level random effects analysis

First, we performed the standard random-effects analysis (e.g., Holmes & Friston, 1998), where individual activation maps are overlaid in the common space, and a *t*-test is performed across participants in each voxel for each relevant contrast. In particular, following DB, we examined group-level effects for the following four contrasts: (i) *Semantic>Fixation;* (ii) *Syntactic>Fixation;* (iii) *Semantic>Syntactic;* and (iv) *Syntactic>Semantic.*

#### 2.4.2. Activation-peak-based, group-level ROI analysis

Next, we performed a more targeted analysis of the activation peaks that emerged in DB for the direct contrasts of the Semantic and Syntactic conditions: for the *Semantic>Syntactic* contrast, the reported peak was at {-48, 20, -4} in Talairach space ({48.5, 20.8, -3.6} in the MNI space); for the *Syntactic>Semantic* contrast, the reported peak was at {-44, -22, 10} ({-44.4, -23.2, 9.7} in the MNI space). To do so, we defined spherical ROIs (of two different sizes: radius = 5mm and 10mm; available for download from https://evlab.mit.edu/papers/Siegelmanbioarxiv) around those activation peaks and extracted the responses to the Semantic and Syntactic conditions (including effects specific to each type of construction). We then performed one-tailed t-tests to evaluate whether the previously reported effects replicate in the current dataset.

#### 2.4.3. Individual-level functional ROI analysis

Finally, we gave the data the strongest chance to reveal a dissociation if such is present, using an individual-subject functional localization approach, which has been shown to benefit from higher sensitivity and functional resolution compared to traditional group-based whole-brain or ROI analyses (e.g., Nieto-Castañon & Fedorenko, 2012). In particular, we used anatomical masks (Tzourio-Mazoyer et al., 2002) for the three subregions of the left inferior frontal gyrus (LIFG) – orbital (LIFGorb), triangular (LIFGtri), and opercular (LIFGop) – and within each of these masks we searched, in each participant individually, for the most Semantics-selective voxels (i.e., showing the strongest effect for the *Semantic>Syntactic* contrast), and for the most Syntax-selective voxels (i.e., showing the strongest effect for the *Syntactic>Semantic* contrast). To define individual functional regions of interest (fROIs), we divided the data in half, and using one half of the data we sorted the voxels within each mask by the t-value for the relevant contrast (i.e., *Semantic>Syntactic* or *Syntactic>Semantic).* We then chose the top 10% of voxels as the fROI. Thus, in each participant, we defined 6 fROIs: (i) a Semantics-selective fROI in LIFGorb; (ii) a Semantics-selective fROI in LIFGtri; (iii) a Semantics-selective fROI in LIFGop; (iv) a Syntax-selective fROI in LIFGorb; (v) a Syntax-selective fROI in LIFGtri; and (vi) a Syntax-selective fROI in LIFGop. We then extracted the responses to the Semantic and Syntactic conditions (including effects specific to each type of construction) from the other half of the data.

This analysis can help circumvent the high inter-individual variability that characterizes the mapping of function onto anatomy in human frontal lobes (e.g., Amunts et al., 1999; Tomaiuolo et al., 1999; Juch et al., 2005; Fedorenko et al., 2012b). Thus, even if the individual activation peaks for the *Semantic>Syntactic* and *Syntactic>Semantic* contrast are spatially variable enough so that group-level analyses (both whole-brain random-effects analysis and ROI-based analysis) fail to detect them, this analysis would recover these effects if they hold across participants anywhere within the LIFG.

The individual activation maps for the *Semantic>Fixation* and *Syntactic>Fixation* contrasts are available for download at https://evlab.mit.edu/papers/Siegelmanbioarxiv.

## 3. Results

### 3.1. Behavioral results

The original study by DB established that the two conditions are comparable in difficulty by collecting behavioral data from the scanned participants in a separate behavioral study (conducted at least 6 months after the fMRI session). We replicate similar across-condition accuracies and reaction times (RTs) in our study (Figure 2), although we collected behavioral data during the scanning session. In particular, the accuracies for both conditions were close to 90% and not significantly different (*t*_(14)_=0.13, Cohen's *d*=0.03, n.s.). Similarly, RTs did not differ either when considering all trials (*t*_(14)_= −0.09, d=−0.02, n.s.) or when considering correctly answered trials only (*t*_(14)_=0.86, d=0.22, n.s.). Comparable behavioral performance suggests that whatever differences might be observed in neural responses between the two conditions would not be attributable to differences in cognitive effort.

**Figure 2.**
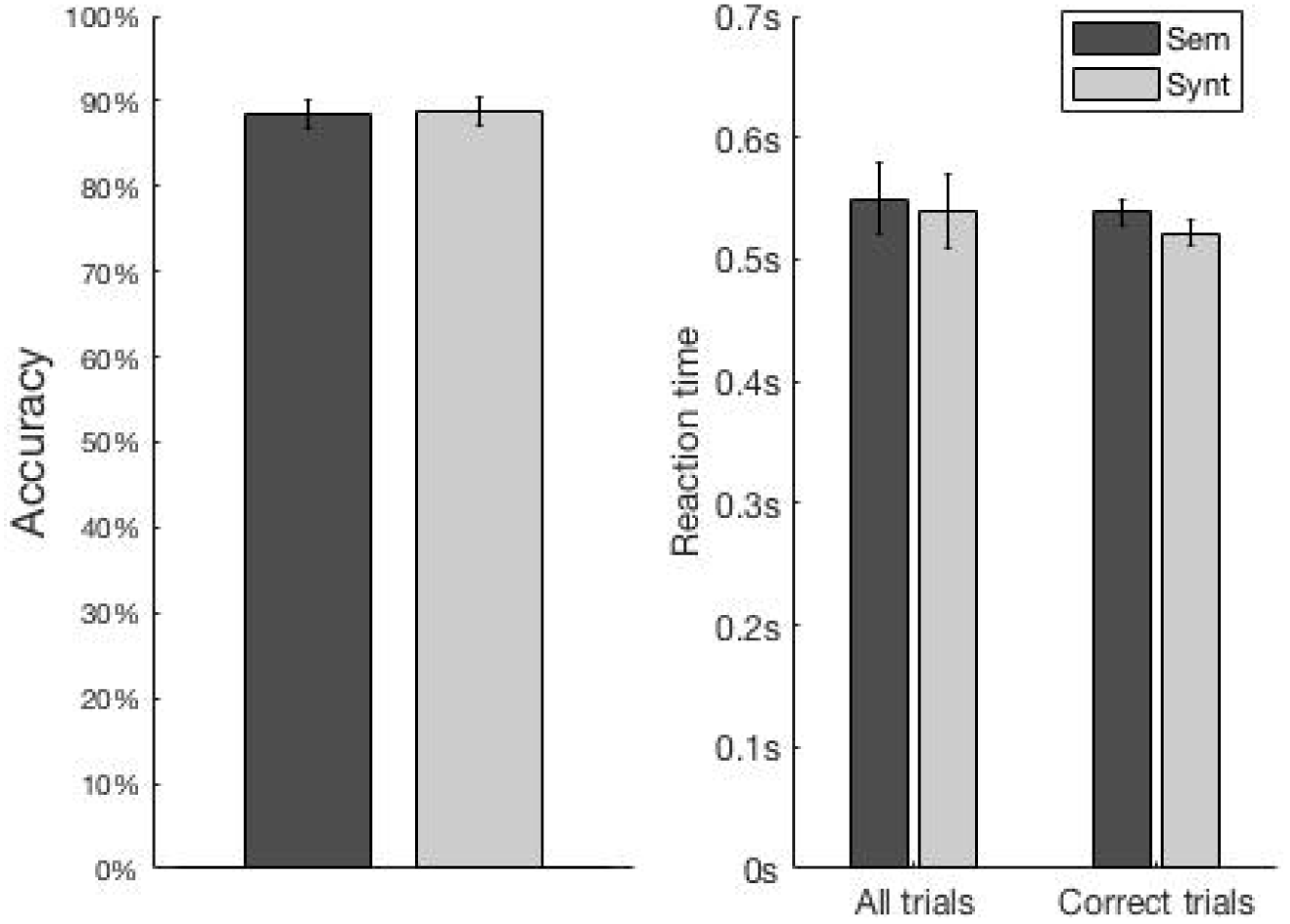
Behavioral performance (left: accuracies; right: RTs) during the Semantic and Syntactic conditions. Error bars represent standard errors of the mean.

### 3.2. fMRI results

#### 3.2.1. Traditional, group-level random-effects analysis

Figure 3 shows whole-brain activation maps for each condition relative to the fixation baseline across the two studies. Visual examination of the maps suggests broad similarity between studies (see also Table SI-1 for a list of the activation peaks for each contrast in the current study), with, critically, robust responses detected for both contrasts in the left inferior frontal cortex.

**Figure 3:**
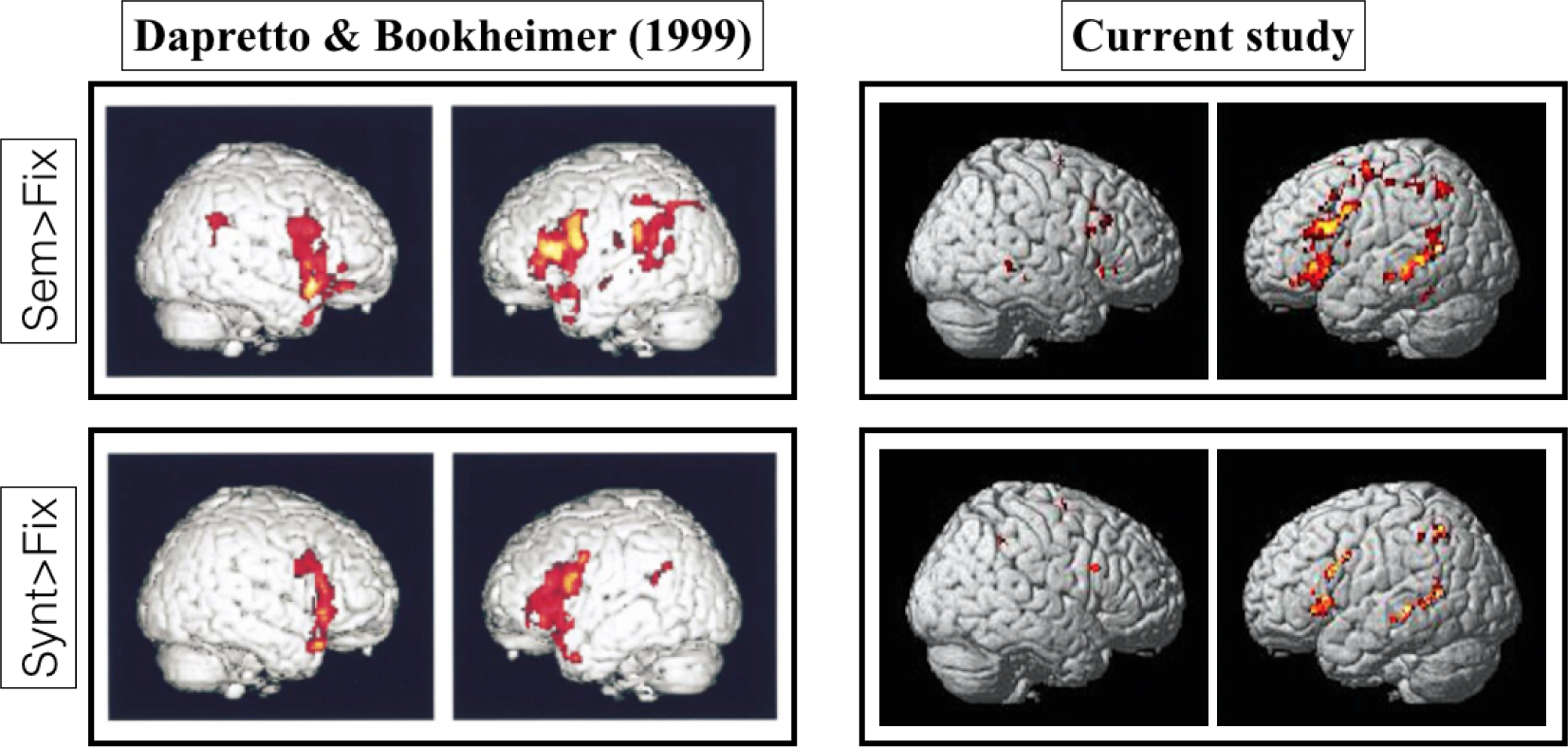
Whole-brain, random-effects activation maps for the *Semantic>Fixation* (top) and for the *Syntactic>Fixation* (bottom) contrasts in DB (left; *p* < 0.0001, uncorrected) and the current study (right; also, *p* < 0.0001, uncorrected; see Table SI-1).

Figure 4 shows the whole-brain activation map for the *Semantic>Syntactic* contrast in DB’s study and our study (centering the crosshair on the same stereotactic location). The *Syntactic>Semantic* contrast did not reveal any significant peaks at thresholds of either 0.0001 or 0.001. The group-level maps for all four contrasts are available at https://evlab.mit.edu/papers/Siegelmanbioarxiv.

**Figure 4:**
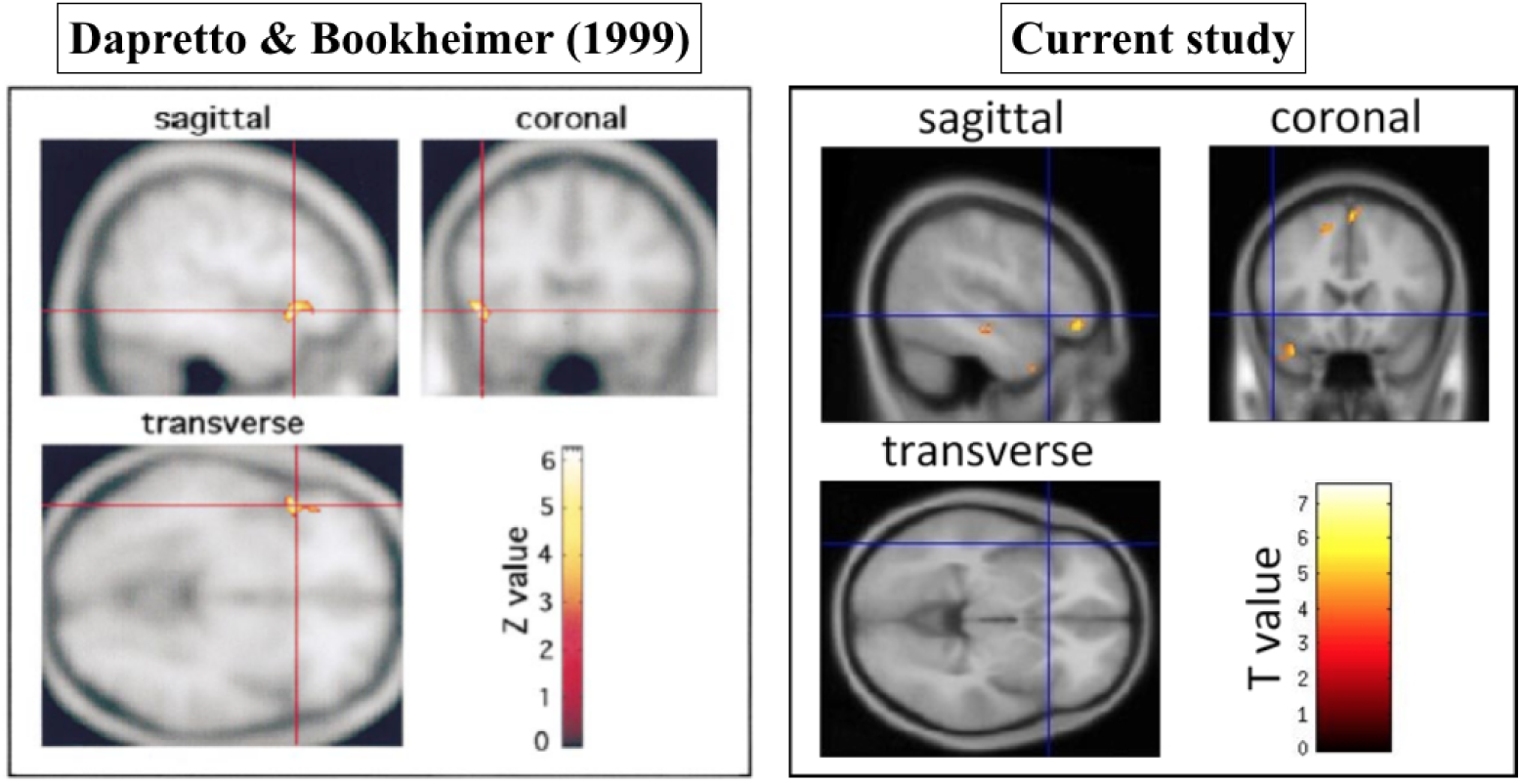
Whole-brain activation maps for the *Semantic>Syntactic* contrast in DB (left; *p* < 0.0001, uncorrected) and the current study (right; also, *p* < 0.0001, uncorrected). The crosshair for the map in the current study was centered on the peak reported in DB for ease of comparison.

#### 3.2.2. Activation-peak-based, group-level ROI analysis

Mean responses to the Semantic and Syntactic conditions in our experiment, including effect specific to each type of construction (Active/Passive *vs.* DO/PP), are shown in Figure 5 for each of the two activation peaks reported in DB. The critical conditions did not show a reliable difference in either the *Semantic>Syntactic* ROI or in the *Syntactic>Semantic* ROI, regardless of the size of the ROI (*ts*< 1.01, *ds*<0.26, *ps*>0.32), plausibly because of the low sensitivity of group-level ROIs (e.g., Nieto-Castanon & Fedorenko, 2012). Numerically, the Semantic condition elicited stronger responses than the Syntactic condition in both ROIs. Further, this pattern was almost identical across the two constructions, providing some degree of generalizability, and arguing against the idea that the failure to replicate the dissociation is due to the changes in the materials (i.e., even when considering the Active/Passive alternation, which was shared between the current study and the original study, the pattern of responses is the same, with the Semantic condition producing a stronger response in both ROIs).

**Figure 5.**
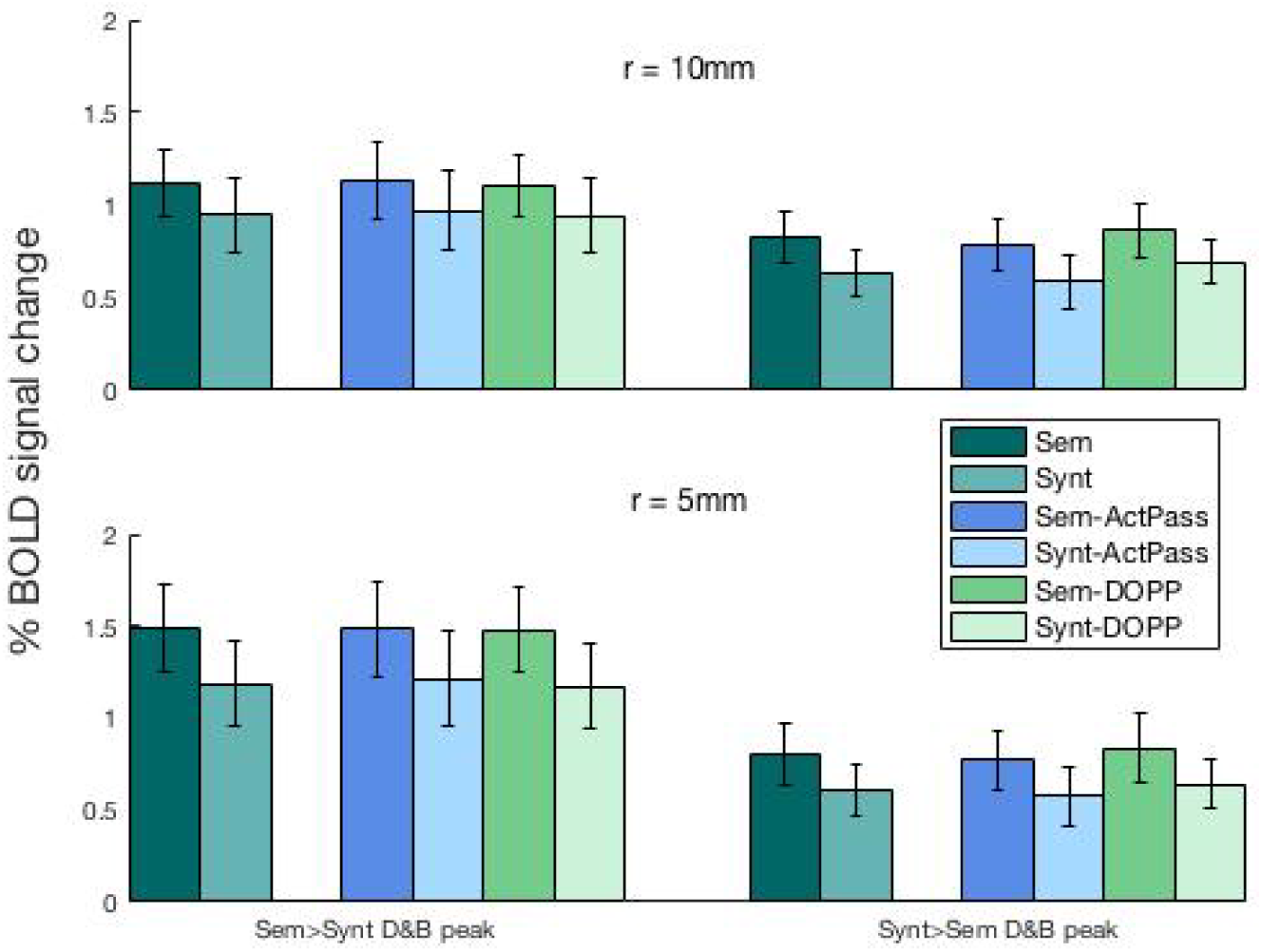
Responses to the critical Semantic (dark teal) and Syntactic (light teal) conditions in ROIs (of two different sizes; 10mm radius – top, and 5mm radius – bottom) defined around the peak coordinates reported by Dapretto & Bookheimer for the *Semantic>Syntactic* contrast, and for the *Syntactic>Semantic* contrast. Blue and green bars further show responses broken down by construction. Error bars represent standard errors of the mean.

#### 3.2.3. Individual-level functional ROI analysis

Figure 6 shows responses to the critical conditions in individually defined functional ROIs. Half of the functional data was used to select the most Semantics-*vs.* Syntax-responsive voxels in each participant separately within each of the three portions of the LIFG, and the responses were then estimated using the other half of the data. This analysis revealed reliable *Semantic>Syntactic* effects in the Semantic>Syntactic fROIs (i.e., fROIs consisting of most Semantics-selective voxels) within LIFGorb (*t*=2.41, *d*=0.55, *p*<0.05), LIFGtri (*t*=2.28, *d*=0.59, *p*<0.05), but not LIFGop (*t*=1.08, *d*=0.28, n.s.).

**Figure 6.**
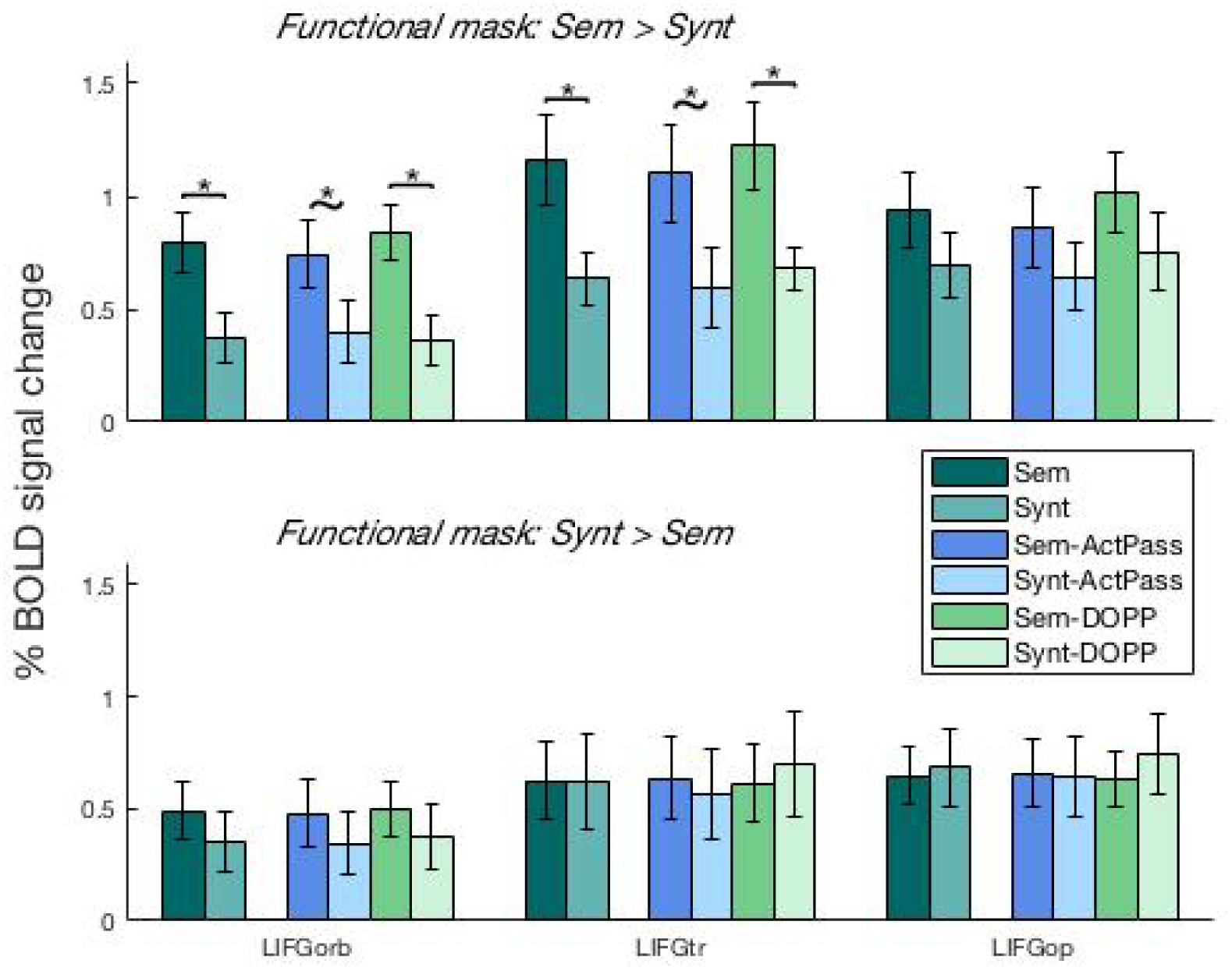
Responses to the critical, Semantic (dark teal) and Syntactic (light teal), conditions in individual fROIs defined by intersecting anatomical masks for the three sub-regions of LIFG (LIFGorb, LIFGtri, and LIFGop) and individual functional maps for the *Semantic>Syntactic* contrast, and for the *Syntactic>Semantic* contrast (in half of the data not used for estimating the responses). Error bars represent standard errors of the mean by participants.

These effects suggest that LIFGorb and LIFGtri contain sub-areas that show robustly and replicably (across runs) greater engagement during the Semantic condition than the Syntactic condition. Furthermore, these effects seem stable across the two constructions: reliable for the DO/PP constructions in LIFGorb (*t*=2.94, *d*=0.76, *p*<0.01) and LIFGtri (*t*=2.49, *d*=0.64, *p*<0.05), and marginal for the Active/Passive constructions in LIFG orb (*t*=1.71, *d*=0.44, *p*<0.1) and LIFGtri (*t*=1.81, *d*=0.47, *p*<0.1).

In contrast, the analysis of Syntactic > Semantic fROIs (i.e., fROIs consisting of most Syntax-selective voxels) did not reveal any replicable *Syntactic>Semantic* effects in any of the three parts of the LIFG (*ts*<1, *ds*<0.26, *ps*>0.46). In fact, within the LIFGorb, the responses still showed a numerically stronger response for the Semantic than Syntactic condition. This is striking given that we specifically searched for most Syntax-selective voxels, and suggests that no voxels within LIFG respond robustly and replicably (across runs) more strongly during the Syntactic condition than the Semantic condition.

## 4. Discussion

In a now classic fMRI study, Dapretto & Bookheimer (1999) reported a dissociation between semantic and syntactic processing within the left inferior frontal gyrus. We here reported an fMRI study designed to replicate this early result. We used the same two-condition design, but substantially expanded the set of experimental materials (five-fold), and included almost twice as many participants in order to increase statistical power. Although the group-level whole-brain maps contrasting each condition to a low-level baseline revealed broad similarity between the two studies (as well as between the semantic and syntactic conditions), the direct contrasts of the Semantic and Syntactic conditions did not replicate the originally reported dissociation. In particular, we found a number of reliable activation peaks for the *Semantic>Syntactic* contrast, including within the LIFG, but the *Syntactic>Semantic* contrast did not produce any reliable peaks within LIFG. In line with this whole-brain analysis, we found a similar pattern in group-level ROIs defined around the *Semantic>Syntactic* and *Syntactic>Semantic* activation peaks reported in DB: the Semantic condition elicited numerically, though not reliably, greater response in both the Semantic-peak ROIs, and the Syntactic-peak ROIs. Finally, in an individual-subjects functional localization analysis, which circumvents the inter-individual anatomical and functional variability that is rampant in the left frontal lobe (e.g., Amunts et al., 1999; Tomaiuolo et al., 1999; Juch et al., 2005; Fedorenko et al., 2012b), we were able to detect reliably greater responses to the Semantic than Syntactic condition within the orbital and triangular portions of the LIFG. However, nowhere within the LIFG were there regions that responded reliably more strongly during the processing of the Syntactic condition compared to the Semantic condition. Thus, the dissociation originally reported in DB does not appear to be robust to replication. (Note that the stronger response to lexico-semantic than syntactic processing in language-responsive cortex is in line with other recent findings, such as the more robust representations of lexical than structural information in the fine-grained patterns of activation, as revealed with multivariate analyses; Fedorenko et al., 2012).

What can explain the non-replication of the original finding? A few factors are likely to be responsible. The first, and perhaps most plausible, contributor is the fact that DB appear to have relied on an analysis that treated participants as fixed effects rather than random effects. In a fixed-effects analysis, individual subjects are not viewed as being randomly drawn from the population. Consequently, the results cannot be generalized beyond the sample tested, and the effects could be potentially driven by a single participant. The seminal publication about this significant limitation in many early brain-imaging studies had only come out a year earlier (Holmes & Friston, 1998), and thus it is possible that the authors had still relied on the fixed-effects analysis (it is difficult to determine this with certainty from the description provided in their Methods section; however, DB do not cite Holmes & Friston (1998), unlike other fMRI studies that were published in the same year and relied on the then-novel random effects analysis).

Second, the original study used a small number of experimental items (8 per condition). This number is very low by the standards of language research, especially in cognitive neuroscience studies, where at least 20-30 items per condition are typically used to ensure generalizability. In the current study we used 40 unique trials per condition, and observed highly similar patterns across two distinct constructions. Thus, it is possible that in the original study, one or two of the items were driving the effects (see e.g., Bedny et al., 2007, for discussion).

It is also worth noting that DB, as is not uncommon in the fMRI literature, did not report the magnitudes of response to the Semantic and Syntactic condition, so that one cannot determine the effect sizes in their study (only their significance is reported). Therefore, the significant peaks reported in DB are consistent with effect sizes anywhere between large and relatively small. Two extreme possibilities consistent with their data are depicted in Figure 7. We suspect that the original result was more consistent with the possibility shown in the right panel of Figure 7, i.e., with small effect sizes; and small effects, especially observed in underpowered studies, are less likely to be real (e.g., Gelman & Carlin, 2014; Simonsohn, 2015; Open Science Collaboration, 2015).

**Figure 7.**
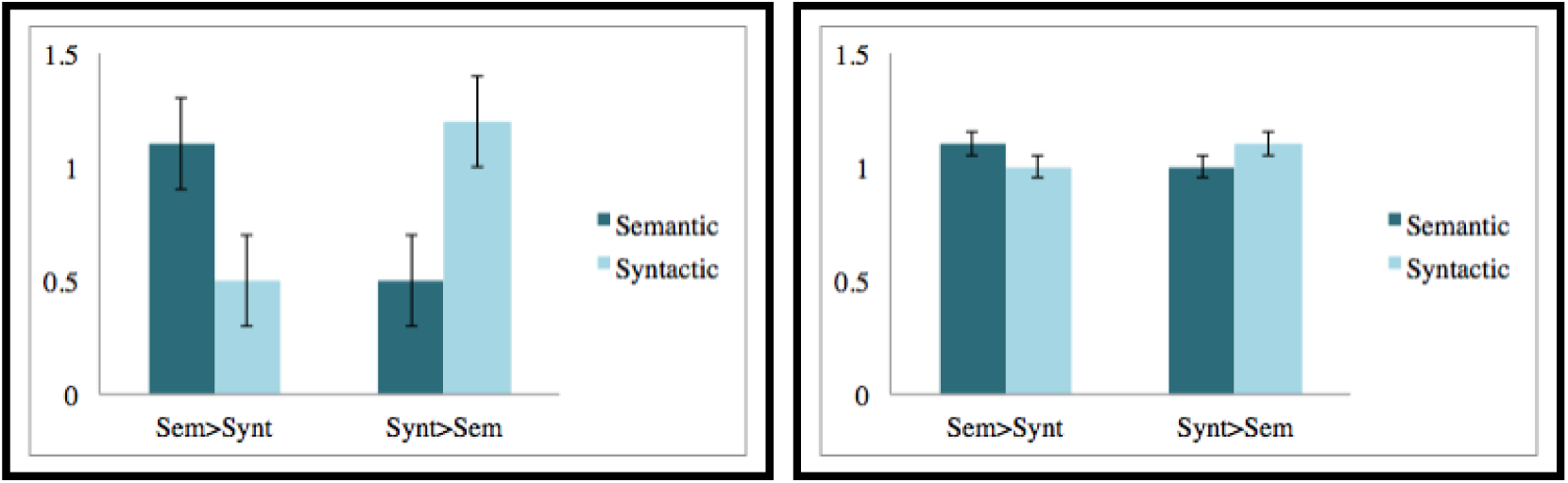
Hypothetical patterns of response in the Dapretto & Bookheimer study, with either large (left) or small (right) effect sizes for the critical contrasts. Conventions are the same as in Figure 5.

To conclude, although the question of whether distinct pools of cognitive resources and cortical regions support lexico-semantic and syntactic processing is likely to keep generating controversy and further research, we here found that at least one study that is commonly cited as evidence for this dissociation does not appear to replicate in a study that employs similar design and materials and has greater statistical power. It may be important to ask, as researchers have recently done in the field of psychology (e.g., Open Science Collaboration, 2015), what proportion of fMRI studies are robust to replication.

## Acknowledgments

E.F. was supported by the NICHD Award HD-057522. We also acknowledge the Athinoula A. Martinos Imaging Center at the McGovern Institute for Brain Research, MIT. For technical support during scanning, the authors thank Atsushi Takahashi and Steve Shannon.

## Supplemental Information

**Table SI-1.** Activation peaks from the group-level, random-effects analysis. The peaks for the Syntactic and Semantic conditions (relative to the fixation baseline) and for the *Semantic>Syntactic* contrast are reported for *p* < 0.0001. The peaks for the *Syntactic>Semantic* contrast are reported for *p* < 0.01.

| Region (Brodmann Area) |  |  | Syntactic>Fixation |  |  |  | Semantic>Fixation |  |  |  |
| --- | --- | --- | --- | --- | --- | --- | --- | --- | --- | --- |
|  |  |  | *x* | *y* | *z* | *t* | *x* | *y* | *z* | *t* |
| Inferior Parietal Lobule | (BA 1) | L |  |  |  |  | -46 | -40 | -58 | 6.06 |
| Postcentral Gyrus | (BA 1) | L |  |  |  |  | -44 | -24 | 54 | 6.30 |
| Middle Frontal Gyrus | (BA 6) | L |  |  |  |  | -24 | -10 | 52 | 7.91 |
| Superior Frontal Gyrus | (BA 6) | L |  |  |  |  | -10 | 8 | 66 | 6.46 |
| Middle Frontal Gyrus | (BA 6) | L |  |  |  |  | -34 | 8 | 46 | 6.43 |
| Middle Frontal Gyrus | (BA 6) | L |  |  |  |  | -40 | 0 | 54 | 5.43 |
| Inferior Parietal Lobule | (BA 7) | L |  |  |  |  | -38 | -44 | 52 | 6.34 |
| Inferior Frontal Gyrus | (BA 8) | L | -42 | 6 | # | 5.69 |  |  |  |  |
| Medial Frontal Gyrus | (BA 8) | L |  |  |  |  | -8 | 22 | 48 | 7.91 |
| Middle Frontal Gyrus | (BA 8) | L |  |  |  |  | -46 | 14 | 46 | 7.57 |
| Middle Frontal Gyrus | (BA 9) | R |  |  |  |  | 52 | 26 | 30 | 6.85 |
| Insula | (BA 13) | L |  |  |  |  | -34 | 24 | 2 | 9.20 |
| Sub-Gyral | (BA 13) | R |  |  |  |  | 32 | 22 | 8 | 6.21 |
| Declive | (BA 19) | R |  |  |  |  | 16 | -64 | -30 | 8.26 |
| Middle Temporal Gyrus | (BA 21) | L | -54 | -46 | 2 | 5.80 | -56 | -34 | -4 | 8.86 |
| Middle Temporal Gyrus | (BA 21) | L | -50 | -26 | -8 | 7.05 | -50 | -24 | -8 | 12.15 |
| Middle Temporal Gyrus | (BA 21) | R | -50 | -34 | 0 | 8.29 | 62 | -40 | 0 | 6.81 |
| Temporal Lobe | (BA 21) | R |  |  |  |  | 50 | -28 | -10 | 5.82 |
| Fusiform Gyrus | (BA 37) | L |  |  |  |  | -36 | -42 | -24 | 7.97 |
| Fusiform Gyrus | (BA 37) | R |  |  |  |  | 38 | -48 | -16 | 6.44 |
| Inferior Parietal Lobe | (BA 39) | L |  |  |  |  | -32 | -58 | 48 | 7.38 |
| Sub-Gyral | (BA 39) | L | -30 | -52 | # | 7.60 |  |  |  |  |
| Superior Temporal Gyrus | (BA 39) | L | -48 | -54 | 8 | 7.25 |  |  |  |  |
| Inferior Frontal Gyrus | (BA 44) | L | -50 | 12 | # | 6.61 |  |  |  |  |
| Inferior Frontal Gyrus | (BA 44) | R | 46 | 14 | # | 5.81 |  |  |  |  |
| Inferior Frontal Gyrus | (BA 47) | L | -34 | 26 | -2 | 8.90 |  |  |  |  |
| Caudate | (BA 48) | R |  |  |  |  | 10 | 8 | 6 | 6.51 |
| Thalamus | (BA 50) | L |  |  |  |  | -12 | -12 | 10 | 7.61 |
| Culmen | undefined | R |  |  |  |  | 2 | -54 | -22 | 6.72 |
| Sub-Gyral | undefined | R |  |  |  |  | 40 | -36 | 4 | 6.72 |
| Extra-Nuclear | undefined | R | 28 | 22 | 0 | 7.63 |  |  |  |  |
| Lentiform Nucleus | undefined | L | -16 | -2 | 8 | 5.96 |  |  |  |  |
| Superior Frontal Gyrus | undefined | L | 0 | 6 | # | 5.81 |  |  |  |  |
| Superior Frontal Gyrus | undefined | L | 0 | 20 | # | 5.49 |  |  |  |  |
| undefined | undefined | L |  |  |  |  | -8 | -58 | -34 | 5.90 |
|  |  |  | Syntactic>Semantic |  |  |  | Semantic>Syntactic |  |  |  |
| Precuneus | (BA 7) | R | # | -68 | # | 3.56 |  |  |  |  |
| Medial Frontal Gyrus | (BA 9) | L |  |  |  |  | -4 | # | 48 | 6.09 |
| Superior Frontal Gyrus | (BA 9) | L |  |  |  |  | -12 | # | 32 | 7.53 |
| Medial Frontal Gyrus | (BA 10) | L |  |  |  |  | -8 | # | 22 | 5.31 |
| Inferior Frontal Gyrus | (BA 47) | L |  |  |  |  | -46 | # | -12 | 6.10 |

